# The creation of sexual dimorphism in Drosophila gonad stem cell niches

**DOI:** 10.1101/367268

**Authors:** Nicole Camara, Cale Whitworth, Mark Van Doren

## Abstract

Sex-specific development of the gonads is a key aspect of sexual dimorphism that is regulated by Doublesex/Mab3 Related Transcription Factors (DMRTs) in diverse animals species. We find that in mutants for *Drosophila dsx*, important components of the male and female gonad stem cell niches (hubs and terminal filaments/cap cells, respectively) still form. Initially, gonads in all *dsx* mutants (both XX and XY) initiate the male program of development, but later half of these gonads switch to form female stem cell niche structures. One individual can have both male-type and female-type gonad niches, however male and female niches are usually not observed in the same gonad, indicating that cells make a “group decision” about which program to follow. We conclude that *dsx* does not act in an instructive manner to regulate male vs. female niche formation, as these structures form in the absence of *dsx* function. Instead, *dsx* acts to “tip the balance” between the male or female programs, which are then executed independent of *dsx*. We show that *bric a brac* acts downstream of *dsx* to control the male vs. female niche decision. These results indicate that, in both flies and mammals, the sexual fate of the somatic gonad is remarkably plastic and is controlled by a combination of autonomous and non-autonomous cues.

## INTRODUCTION

The creation of sexual dimorphism, the differences between the sexes, is a critical aspect of development. Nowhere is this process more important than in the gonads, which must produce the male and female gametes for sexual reproduction, and which often also control the sexual development of other cell types in the organism. A key part of sexual dimorphism in the gonad is the formation of germline stem cells, and the microenvironments or “niches” that control them. In many organisms, both the testis and the ovary contain stem cells, but these systems are different in their morphology and regulation. In other species, such as humans, sexually dimorphic development of the gonads results in very different stem cell potential in the two sexes, with only the testes having a clear stem cell population.

Although animal species vary widely in the mechanisms that trigger sexual identity, downstream components that control sex-specific development may be more well-conserved. The Doublesex/Mab3 Related Transcription Factors (DMRTs) control gonad sexual dimorphism in a wide range of animals, including flies, fish, frogs, birds, mice and man (Matson and Zarkower, 2012). The founding member of this family, *Drosophila doublesex (dsx)*, controls many aspects of sex-specific development (Hildreth, 1965; Baker and Ridge, 1980), including all known sex differences in the somatic gonad (DeFalco et al., 2003; Le Bras and Van Doren, 2006; DeFalco et al., 2008). However, many questions remain about how this conserved class of transcription factors regulates sexual dimorphism and the downstream targets through which they act. Do these transcription factors regulate many targets to “micro-manage” sexual development, or do they instead regulate a few genes that initiate independent developmental pathways? What are the critical time points in development when these factors provide information about sexual identity? Do factors like DSX, which control sexual dimorphism in several different tissues, do so by regulating similar or distinct target genes in these tissues? Here we address some of these fundamental questions for how DSX regulates male vs. female gonad stem cell niche development.

Drosophila sex determination is regulated by X chromosome number (XX is female and XY is male), and the presence of two X chromosomes activates expression of the Sex lethal (SXL) protein (reviewed in (Camara et al., 2008)). SXL initiates an alternative splicing cascade acting through Transformer (TRA) and Transformer 2 (TRA2) to regulate RNA splicing of *dsx* and *fruitless (fru)*, which encode the key transcription factors controlling sexual dimorphism. *fru* is thought to act primarily in the nervous system to influence sexual behavior (reviewed in (Villella and Hall, 2008),) while *dsx* influences behavior as well as most other aspects of sex-specific morphology (reviewed in (Camara et al., 2008; Dauwalder, 2011)). While only the male splice-form of *fru* produces a functional protein, alternative splicing of *dsx* produces functional isoforms in males and females (DSX^M^ and DSX^F^, respectively) (Baker and Wolfner, 1988; Burtis and Baker, 1989). DSX^M^ and DSX^F^ share a common Zn-finger DNA binding domain but have different C-termini, which confer the ability of these proteins to have different effects on target gene expression. The most well-characterized target, the Yolk Protein gene locus, is activated in the fat body by DSX^F^ and repressed by DSX^M^ (Coschigano and Wensink, 1993). More recently, a few other DSX targets have been identified, including the *bric-a-brac* locus *(bab1* and *2)* which regulates sex-specific abdominal pigmentation (Williams et al., 2008). In addition, genomic studies have predicted many more DSX targets (Chatterjee et al., 2011, Luo et al., 2011, Clough et al., 2014, Arbeitman et al., 2016)

In *Drosophila*, both the ovary and testis have germline stem cells that are controlled and maintained by surrounding somatic cells. A key component of the male stem cell niche is created by the ‘hub’, a tight cluster of cells at the anterior tip of the testis (Aboïm, 1945; Hardy et al., 1979; Kiger et al., 2000; Tulina and Matunis, 2001), which forms during the last stages of embryogenesis (Stage 17) (Gönczy et al., 1992; Le Bras and Van Doren, 2006). In the ovary, each of the roughly 16 ovarioles contains a stem cell niche, key components of which are the cap cells (CC) and terminal filaments (TF) (Xie and Spradling, 2000) (reviewed by (Spradling et al., 1997; Chen et al., 2011)). The female niche develops much later than the male niche, with terminal filaments forming in the late 3^rd^ instar larval period and cap cells forming at the larval to pupal transition (King, 1970; Zhu and Xie, 2003). Although the two niches are quite different morphologically, there are similarities in how they act to nurture the germline stem cells (Gilboa and Lehmann, 2004; Fuller and Spradling, 2007; Dansereau and Lasko, 2008).

Here we study the role of *dsx* in controlling the development of the male and female stem cell niches. We find that important components of the niche, the hub in males and the CC/TF in females, can form in the absence of *dsx* function, but do so stochastically in both XX and XY individuals. Thus, *dsx* is not required to instruct cells how to form these structures, but is instead only required to ensure that the proper structures form in the correct sex. We propose that *dsx* activates endogenous pathways for male and female niche formation which then act independently of *dsx* function, and we identify *bric-a-brac* as a downstream target by which *dsx* can activate such a pathway. We also find that, while the hub forms initially in all *dsx* mutants, half of the gonads (both XX and XY) lose the hub and form CC and TF, apparently from some of the same cells that initially formed the hub. Thus, the gonad stem cell niches are remarkably plastic in their developmental programs. We have been able to determine the critical time points for this developmental plasticity using a conditional allele of the sex determination gene *tra2*. Lastly, we find that male and female niches do not form within the same gonad indicating that cells communicate with one another about whether to follow the male or female program.

## RESULTS

### *dsx* mutant adult gonads have either a male-like or female-like stem cell niche

At the end of embryogenesis male and female gonads are already different, as evidenced by the presence of three male-specific cell types: male-specific somatic gonadal precursors (msSGPs), pigment cell precursors (PCPs) and hub cells (DeFalco et al., 2003; Le Bras and Van Doren, 2006; DeFalco et al., 2008). In previous work, we examined the fate of each of these cell types in *dsx* mutants and found that, in the absence of *dsx*, msSGPs, PCPs and hub cells are present in both XX and XY *dsx* mutant embryonic gonads (DeFalco et al., 2003; Le Bras and Van Doren, 2006; DeFalco et al., 2008). Thus, embryonic gonad development in *dsx* mutants begins along a male pathway, and *dsx* is not required for the initial formation of any of these cell types.

We wanted to assess what happens to the hubs in *dsx* mutants later in development. In wild type (wt), XX animals produce TF/CC (Fig. 1A) while XY animals always form hubs (Fig. 1B). Surprisingly, we found that in both XX and XY *dsx* mutant adults, half of the gonads had a hub, characteristic of a male gonadal niche, and the other half had terminal filaments (TFs), characteristic of a female gonadal niche (XX: 48% hub, 52% TF n=144; XY: 48% hub, 52% TF n=104) (Fig. 1C-F). To assess hub cell identity, we examined expression of three known hub markers; N-Cadherin, Fascilin3 (Fas3), and an *escargot (esg)* enhancer trap which expresses in hub cells, *(esg^M5-4^)* (Gönczy and DiNardo, 1996). All three markers are expressed in *dsx* mutant hubs (Figs 1C-G, S1A and data not shown). To assess TF identity, we used two known markers for TFs, Sox100B (Nanda et al., 2009) and Engrailed (EN, (Forbes et al., 1996)), and found that they are expressed in *dsx* mutant TFs (Figs 1H, S1B). In addition to TFs, cap cells are an important part of the female stem cell niche. We assessed whether cap cells were present in gonads with TFs using a co–stain for LaminC (LamC) and Zinc finger homeodomain 1 (Zfh-1). LaminC is expressed strongly in the terminal filament and weakly in cap cells (Xie and Spradling, 2000), and Zfh-1 is expressed weakly in terminal filaments and strongly in cap cells. We found that those *dsx* mutants which had TFs present also had cap cells (Figs 1I, S1E). We also assessed cap cells and TFs using an enhancer trap in *hedgehog* expressed in these cells (Forbes et al., 1996), which also indicated that both of these cell types are present in *dsx* mutants (Fig. S1C,D). Some of the molecular markers used are expressed in both hubs and TFs (e.g. N-Cadherin, *hh-lacZ*), but the clear differences in hub vs. TF morphology (hubs are compact clusters of cells, while TF are linear chains, testes have a single hub while TF are found in each of the 16 or so ovarioles per ovary) are a strong indication of the hub vs. TF developmental program and identity. In addition, some molecular markers are specific for hubs (Fas3) or TFs (lamC and Sox100B). These markers were always expressed in the appropriate structure in *dsx* mutants, regardless of whether the animal was XX or XY; Fas3 was on in hubs but not TF (e.g. Fig. S4C) and Sox100B and lamC were always on in TF but not hubs (e.g. Fig. 5A-D).

**Fig 1.**
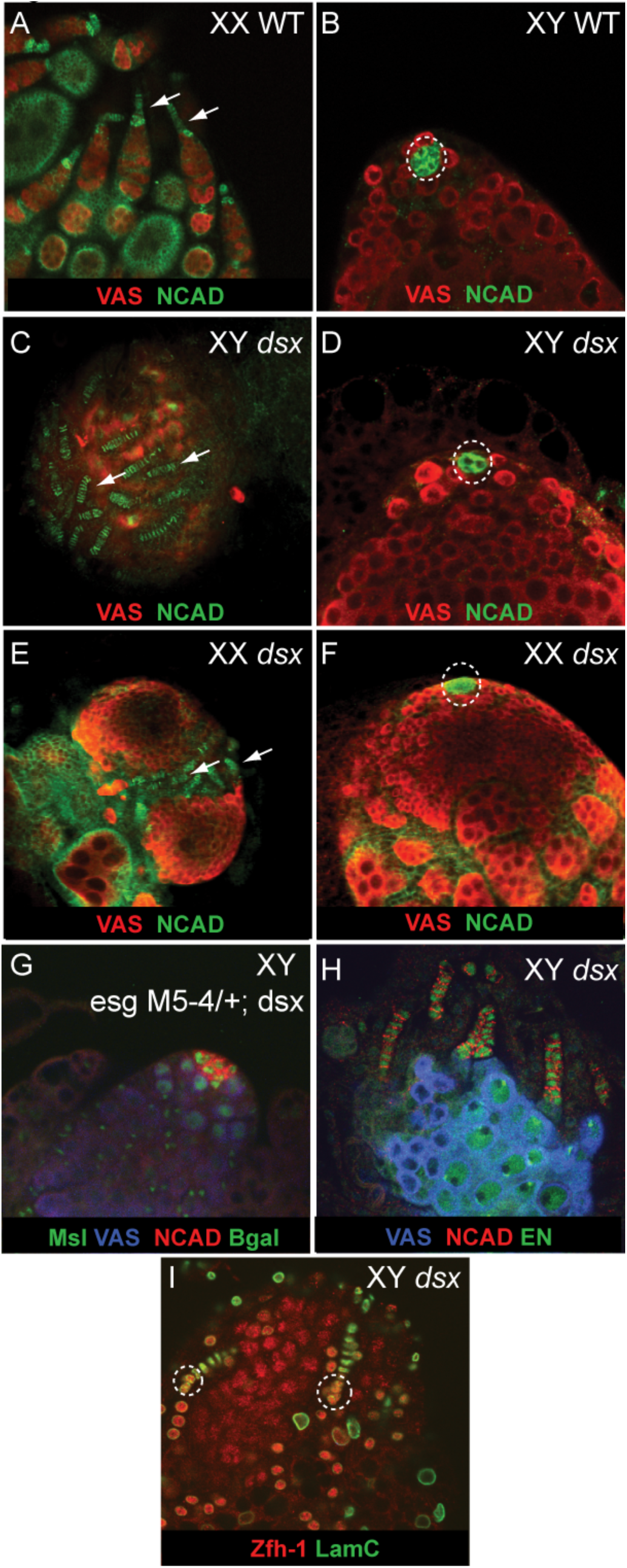
*dsx* mutant adults have either hubs or terminal filaments. Staining as indicated in all panels. (A, B) Wild type adult gonads. (Terminal filaments, arrows; hubs, circled) (C-I) *dsx* mutant adult gonads. (C,E) Terminal filaments in XY and XX *dsx* mutants. (D, F) Hubs in XY and XX *dsx* mutant gonads. (G) *esg* enhancer trap expression in *dsx* mutant hub. MSL= anti-MSL2 to determine sex chromosome genotype. (H) Engrailed (EN) expression in *dsx* mutant TFs (I) Cap cells (Labeled with anti-Zfh-1 and anti-LamC) in *dsx* mutant gonads with TFs.

In order to determine whether the hubs in *dsx* mutants are able to properly signal to the surrounding stem cells, we examined activation of STAT92E in germ cells adjacent to the hub, which can be used as an assay for activation of the JAK/STAT pathway (Wawersik et al., 2005). Germ cells adjacent to hubs in *dsx* mutants exhibited increased STAT92E immunoreactivity, similar to GSCs in wild-type (wt), indicating that hubs in *dsx* mutants are functional (Figs 2B, S2A), although STAT92E in XX germ cells appeared less intense than in XY germ cells (Sheryl Southard, Pradeep Bhaskar and MVD, in preparation). We next tested whether gonads exhibiting TFs and CC were capable of signaling to female GSCs by examining an enhancer trap in *Daughters against dpp (Dad-lacZ)*, which is activated in response to TGF-β signaling in the female niche (Casanueva and Ferguson, 2004). At least some germ cells adjacent to a female-like niche expressed *Dad-lacZ*, suggesting the female-like niche in *dsx* mutants is functional (Fig S1F).

**Fig 2.**
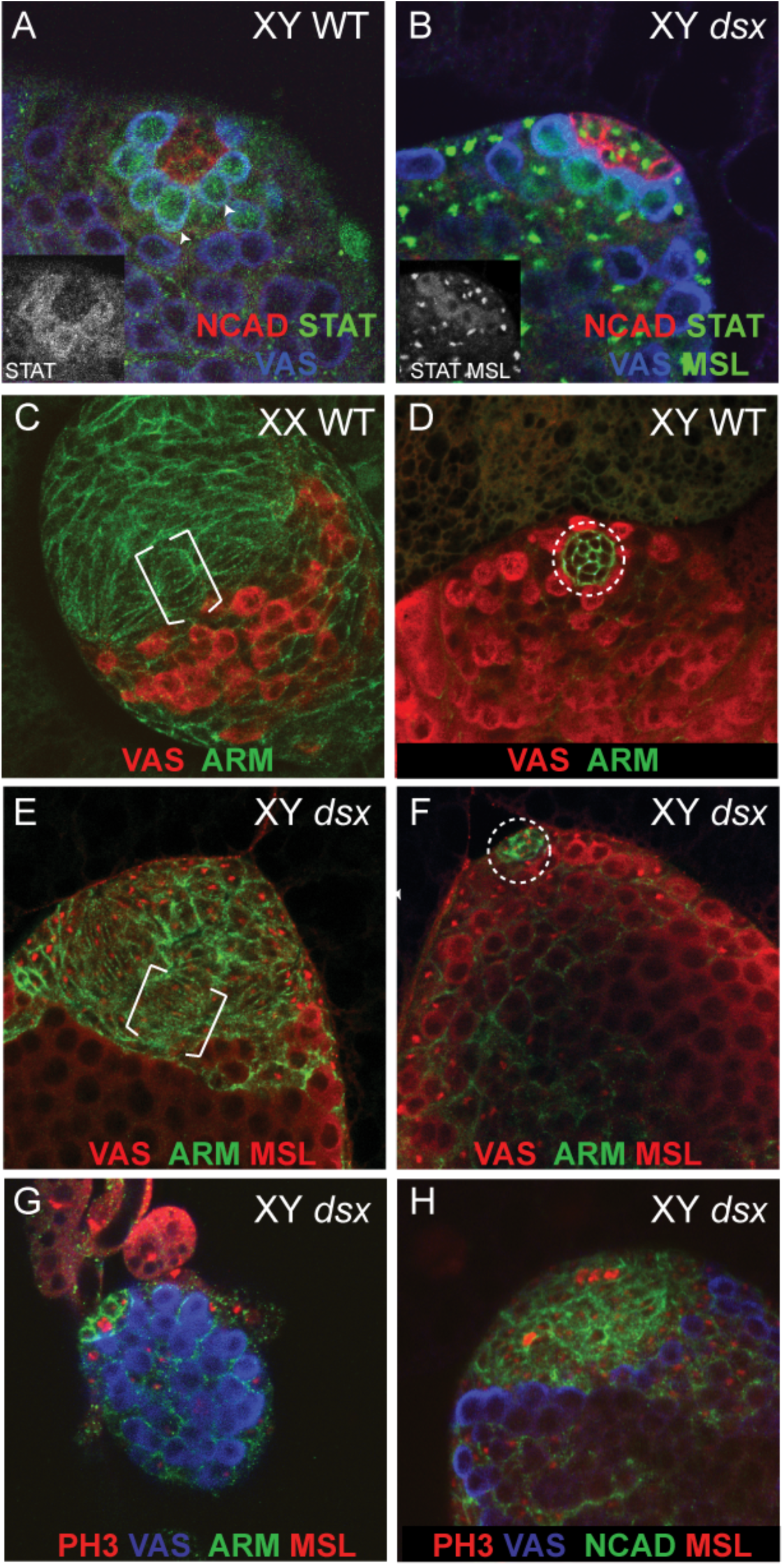
*dsx* mutant niches during the larval stages. Staining as indicated in all panels. (A) STAT92E localization in wild type male germ cells adjacent to the hub. (B) STAT92E localization in *dsx* mutant germ cells adjacent to the hub. Inset shows STAT and MSL channels alone. STAT is cytoplasmic staining in germ cells, MSL is punctuate in XY somatic cells. (C,D) WT niche formation in 3^rd^ instars. Developing terminal filaments are marked with brackets, hubs circled. (E) Developing terminal filaments in XY *dsx* mutant 3^rd^ instar gonad. (F) Hub in *dsx* mutant 3^rd^ instar. (G) Proliferation of hub cells in *dsx* mutant 2^nd^ instar gonad. (H) Proliferation of hub cells in *dsx* mutant 3^rd^ instar gonad.

We conclude that, in the absence of *dsx* function, either a male-like or female-like niche forms, as opposed to formation of an intersexual niche structure. This appears to be a stochastic decision, as approximately 50% of gonads form male or female niches, regardless of the chromosomal constitution (XX vs XY). Thus, *dsx* is not required for the formation of these niches, but is required to ensure that the proper niche forms in the proper sex. Further, in *dsx* mutants, either a hub or TF could be identified in a particular gonad, but rarely both, suggesting that the cells of the gonad are making a group decision as to whether to form hubs or TFs. However, this is not an organism-wide decision, as the two gonads from one individual can differ in whether they have a hub or TFs (paired stainings, data not shown). Finally, in XX *dsx* mutant gonads, pseudo-egg chamber-like structures were found in gonads containing terminal filaments and in gonads containing hubs (Fig 1E,F), but these structures were never observed in XY *dsx* gonads. This suggests that the germ cells in *dsx* mutants partly retain their sexual identity, and that they influence the identity and behavior of surrounding somatic cells. Alternatively, there could be dsx-independent aspects of somatic sexual identity in follicle cell specification.

### Hubs in *dsx* mutants appear to give rise to terminal filaments

Since all *dsx* mutant embryos form hubs, but at adult stages half of the gonads have hubs and the other half have TFs, we first wanted to determine when gonads transition from having male niches to having female niches. Although hubs form by the end of embryogenesis, TF and cap cells do not develop until the late 3rd instar larval period (King, 1970; Zhu and Xie, 2003)(Fig 2C, brackets). In *dsx* mutant 3^rd^ instar larvae, half of the gonads had hubs (Fig 2F and S2B) while the other half instead showed an increased number of somatic cells at one pole of the gonad that were aligning into the “stacks” of cells typical of developing TFs (Figs 2E and S2C). A similar result was observed for both XY and XX *dsx* mutants (XX *dsx* 46% hubs, 54% TFs n=78; XY *dsx* 52.5% hubs, 47.5% TFs n=99) (Figs 2E,F, S2B,C).

Next we wanted to determine how *dsx* mutant gonads transition from hubs to TF. One possibility is that hub cells die while TFs form from a distinct population of cells. However, we found no evidence for hub cell death (anti-capsase 3, data not shown). Another possibility is that the hub cells are trans-fating in some gonads to form TFs. In this case we should observe proliferation of hub cells to give rise to the larger field of TF cells. Normally, proliferation is not observed in larval or adult hub cells ((Hardy et al., 1979) and data not shown). However, in 2nd instar XX and XY *dsx* mutants, we found that hub cells in some gonads were positive for the mitotic marker phospho-Histone H3 (pH3) (Fig. 2G, S2D). In 3^rd^ instar *dsx* mutants that appeared to be transitioning from hubs to TFs, we observed pH3-positive somatic cells in the region where TFs form (Fig 2H, S2E). However, in gonads that appeared to be retaining their hubs, no pH3-positive cells were observed in the hub. These data support the hypothesis that, when gonads switch from hubs to TFs in *dsx* mutants, the same cells that initially form hubs are able to switch their developmental program and proliferate to contribute to TFs. Unfortunately, our attempts to conduct direct lineage analysis of hub cells as gonads transition from hubs to TF were inconclusive for technical reasons.

### Initial somatic gonad identity in *dsx* mutants is intersexual

Our observations indicated that gonad development in *dsx* mutants initiates along a male pathway, but is more plastic than in wt, with half of both XX and XY gonads switching to a female pathway, at least in terms of hubs vs. TF/CC. We reasoned that this could indicate that SGP identity is initially male in *dsx* mutants. Alternatively, SGPs might have an intersexual identity, which is nonetheless sufficient to induce the male pattern of development, but is subject to switching to the female pathway at the time in development that female gonad development begins (during the larval period). To investigate this, we examined SGP identity during gonad formation, prior to hub formation. One sex-specific characteristic of early SGPs is that male SGPs induce increased STAT92E immunoreactivity in all germ cells, indicating that they signal to the germ cells via the JAK/STAT pathway, prior to the time that this response is restricted to male GSCs associated with the hub (Wawersik et al., 2005). We found that in both XY and XX *dsx* mutants, embryonic germ cells expressed STAT92E indicating that SGPs have at least a partial male identity as they can still signal to the germ cells (Fig. 3A,B).

**Fig 3.**
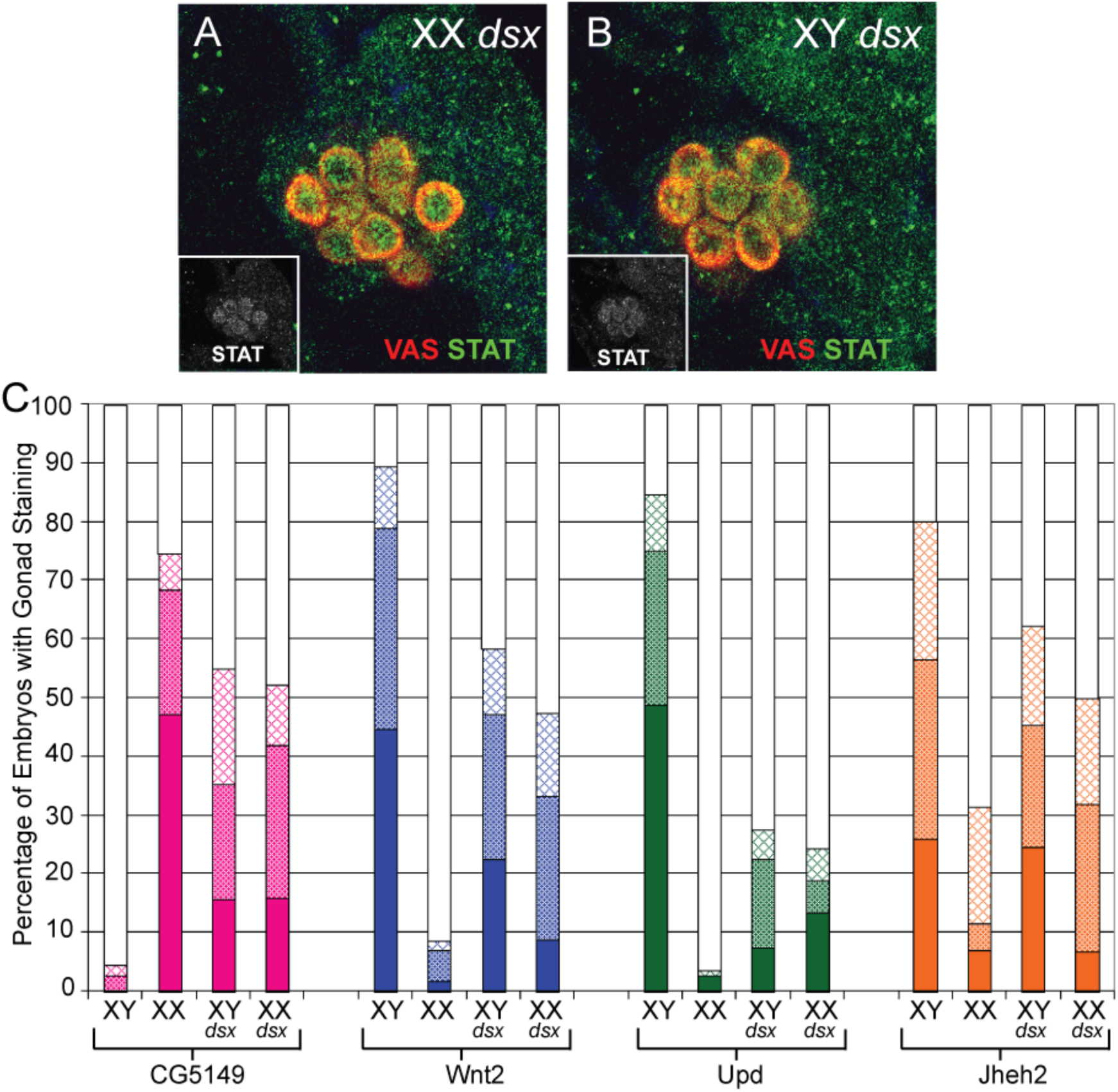
Sexual identity of *dsx* mutant SGPs. Staining as indicated in all panels. (A,B) STAT92E activation in XX and XY *dsx* mutant embryonic germ cells. Inset is STAT channel alone. (C) Graph of sex specific gene expression. Percent of embryos with gonad staining is indicated with bars reflecting intensity of staining (hatched = weak, stippled = medium, solid = strong).

We next analyzed expression of four genes expressed sex-specifically in the SGPs; one female specific gene, *CG5149* (Casper and Van Doren, 2009), and three male-specific genes *Wnt2* (DeFalco et al., 2008), *unpaired* (*upd*) (Wawersik et al., 2005), and *Juvenile Hormone Epoxide Hydrolase 2* (*Jheh2*) (Casper and Van Doren, 2009). We found that in both XX and XY *dsx* mutants, each gene was expressed intermediately between that of a wild-type male and wild-type female (Fig. 3C). Thus, although the gonad begins along a male pathway as evidenced by the presence of msSGPs, hub cells and pigment cells, the sexual identity of the SGPs is not fully masculinized, and retains some mixed character. Apparently, this mixed character is sufficient to initiate the male pathway in the embryo, but the structures formed are more plastic and subject to switching to the female pathway during larval stages.

### Timing of *dsx* action in niche formation

We next wanted to determine when the somatic gonad was competent to transition between hubs and TFs. We used temperature sensitive alleles of *tra2* (*tra2^ts1^/tra2^ts2^*, hereafter referred to as *tra2^ts^*), which in XX animals promotes female development at the permissive temperature (18°C) but male development at restrictive temperature (29°C) (Belote and Baker, 1982). We found that XX *tra2^ts^* animals raised at permissive temperature develop a normal female niche and are fertile, and XX *tra2^ts^* animals develop a hub when raised at restrictive temperature (Fig. 4A,B) (but are sterile due to the XX constitution of the germ cells). An important difference between this experiment and experiments using *dsx* null mutants (above) that lack *dsx* function, is that *tra2^ts^* allows for DSX^M^ expression at the restrictive temperature and DSX^F^ expression at the permissive temperature, and so the cells of the somatic gonad may have a more robust male or female identity, accordingly.

**Fig 4.**
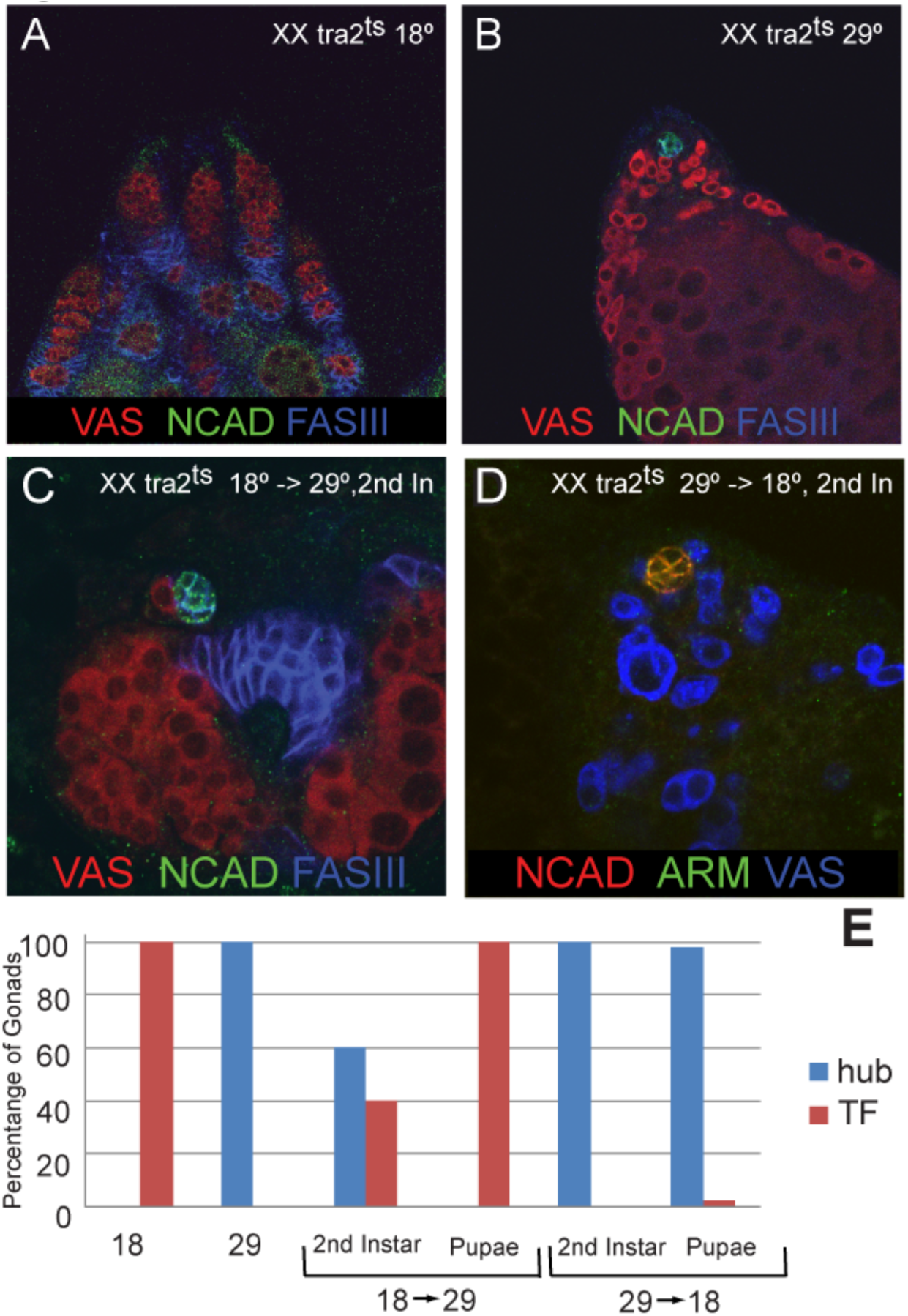
Timing of *dsx* action in niche formation. Staining as indicated in all panels. Note that anti-Fas3 labels hubs but not TF. (A) XX tra2^ts^ animals raised at 18 develop TF. (B) XX tra2^ts^ animals raised at 29 develop hubs. (C) 60% of XX tra2^ts^ animals shifted from 18 to 29 at 2^nd^ instar develop hubs. (D) 100% of XX tra2^ts^ animals shifted from 29 to 18 at 2^nd^ instar maintain hubs. (E) Graph of percentage of XX tra2^ts^ gonads with either hubs (blue bar) or TFs (red bar) in different temperature conditions.

We first raised XX *tra2^ts^* animals at the permissive (female) temperature and then shifted to the restrictive temperature at either 2^nd^ instar (after normal hub formation) or pupal stages (after normal TF formation). When animals were shifted at the second instar stage, 60% of the adult gonads had hubs and 40% had terminal filaments (n=83) (Fig. 4C,E). Thus, even when the soma began development under the influence of DSX^F^, expression of DSX^M^ at the second instar stage was still sufficient to cause hub development, even though this is after the time that hubs would normally form. However, when the same shift was done at the pupal stage, 100% of the gonads had terminal filaments (n=46) (Figs 4E, S3A), suggesting that the sex of the female gonadal niche had been determined and could no longer be influenced by *dsx*.

We performed the reverse experiment by switching XX *tra2^ts^* animals from restrictive (male) to permissive conditions, either at the second instar or pupal stages. When animals were switched at 2nd instar, very little to no germline was present in adults, and a niche could not be identified. Therefore, we examined 3^rd^ instar larval gonads from these animals and found that 100% had hubs (n=38). When the temperature shift was done at the pupal stages, 98% of the gonads formed hubs and 2% (i.e. 1 example) formed terminal filaments (n=49) (Figs 4E, S3B). Thus, hub cells which are fully masculinized under the presence of DSX^M^ are resistant to form TFs, even upon expression of DSX^F^. This supports the conclusion that niches formed in the absence of *dsx* function (*dsx* mutants above) are more plastic, while niches formed in the presence of DSX^M^ or DSX^F^ are more robust. Further, the male and female niches show differences in when their sexual phenotype becomes irreversible, and this timing correlates with when their niches normally form. DSX^M^ acts prior to second instar to irreversibly determine hub fate, while female SGPs are still plastic until late larval stages before DSX^F^ irreversibly determines TF fate.

### Role of *dsx* in msSGP and pigment cell development

In addition to forming hubs, all *dsx* mutant embryonic gonads also develop other male characteristics, including the presence of the msSGPs and pigment cell precursors (DeFalco et al., 2003; DeFalco et al., 2008), and we investigated the role of *dsx* in the later development of these cell types. Although msSGPs express DSX (Hempel and Oliver, 2007), it is not required for their initial development in the embryo (DeFalco et al., 2003; DeFalco et al., 2008). The msSGPs give rise to the terminal epithelium of the testis (Nanda et al., 2009), where the gonad contacts the reproductive tract and the late stages of spermatogenesis occur (Tokuyasu, 1974). In 3^rd^ instar larvae, we found that the terminal epithelium was still present in all XX and XY *dsx* mutant gonads (Fig 5A,B). Unfortunately, we were unable to assay the terminal epithelium in adult *dsx* mutants due to the difficulty of identifying these cells in disorganized gonads without unambiguous molecular markers. Therefore, we conclude that *dsx* is not required for terminal epithelium formation or development up until the 3^rd^ instar larval stage, but we cannot rule out a later role for *dsx* in these cells.

**Fig 5.**
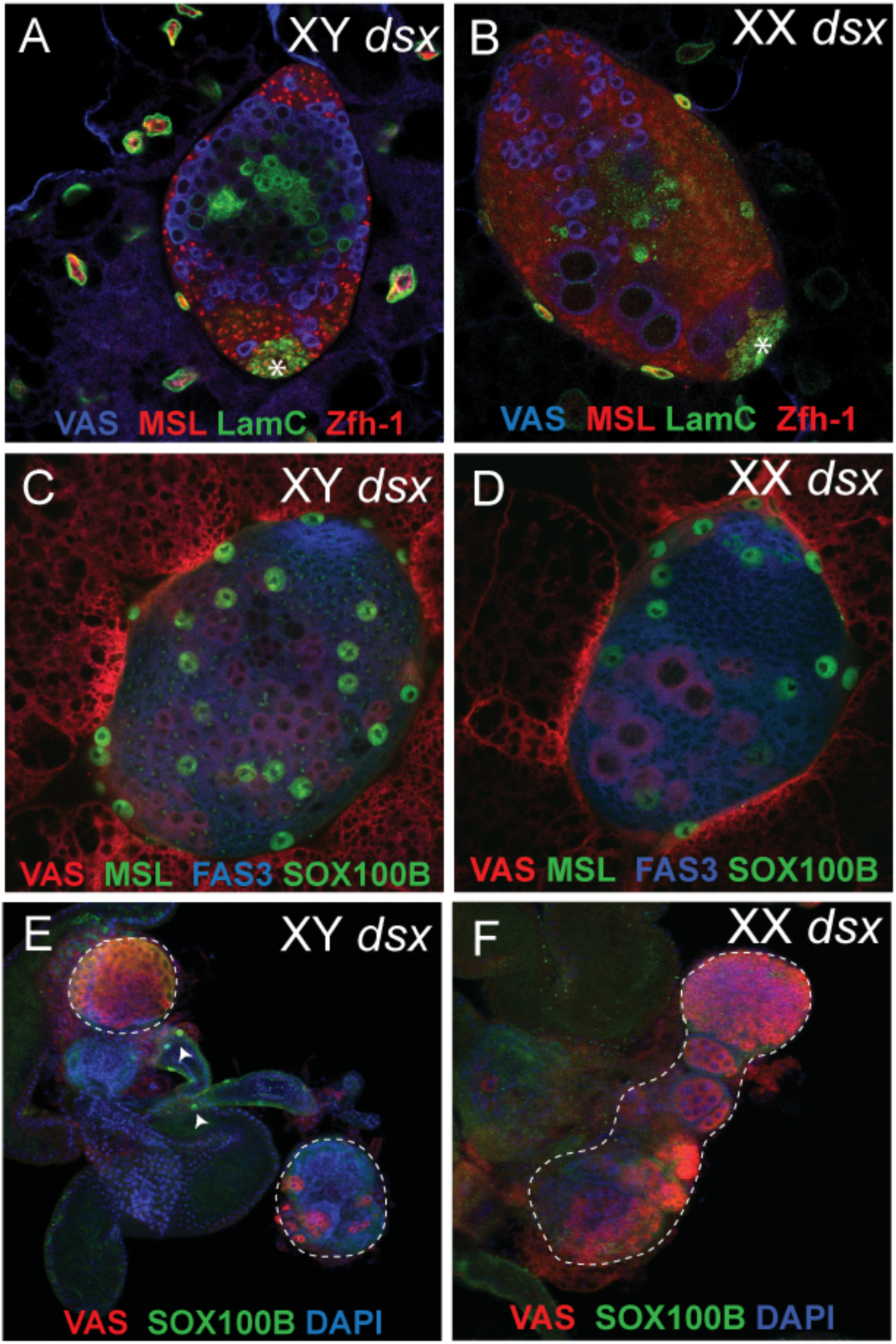
Pigment cell and terminal epithelium development in *dsx* mutants. Staining as indicated in each panel. (A,B) XX and XY *dsx* mutants maintain terminal epithelia. (C,D) XX and XY *dsx* mutants at 3^rd^ instar have pigment cells (large Sox100B positive nuclei). (E,F) XX and XY *dsx* mutants gonads (outlined) have lost their pigment cells by adult stages. However, the seminal vesicle retains pigment cells (arrowheads, Eanimals ).

We next examined the pigment cell precursors, which form a pigmented epithelium around the adult testis (DeFalco et al., 2008). In 3^rd^ instar larvae, pigment cells were observed in both XX and XY *dsx* mutant gonads (Fig 5C,D), but we did not find these cells around the adult gonad (Fig 5E,F). Pigment cells were still present on the seminal vesicle which is a part of the reproductive tract derived from the genital disc (Fig. 5E, arrowheads). DSX is not expressed in the pigment cells themselves (Hempel and Oliver, 2007), and the formation of these cells from the fat body is regulated by male-specific expression of *Wnt2* in the somatic gonad and male genital disc (Kozopas et al., 1998; DeFalco et al., 2008). In *dsx* mutants, Wnt2 expression is reduced in the embryonic gonad (DeFalco et al., 2008) and, though this is sufficient to allow pigment cell precursors to form in the embryo, it may not be enough to maintain the pigment cells after 3^rd^ instar. Alternatively, *Wnt2* expression may be further reduced after 3^rd^ instar in *dsx* mutants. In contrast, the male primordium of the genital disc still develops in *dsx* mutants (Hildreth, 1965), which likely accounts for the maintenance of pigment cells around the seminal vesicle.

### *bab* acts downstream of DSX in sexually dimorphic niche formation

In order to investigate what downstream targets *dsx* might control to regulate sexually dimorphic niche formation, we took a candidate gene approach to find genes that interact with *dsx* in niche formation. The *bric-a-brac 1* and *2 (bab1/2)* locus encodes two related transcription factors and is known to be involved in development of a number of sexually dimorphic tissues, including the abdomen, sex combs and the terminal filaments (Godt et al., 1993; Godt and Laski, 1995; Sahut-Barnola et al., 1995; Kopp et al., 2000; Couderc et al., 2002). We reasoned that if *bab1/2* is a downstream target of *dsx* in controlling male vs. female niche formation, that loss of *bab1/2* function might not only affect TF/CC formation, but might also “tip the balance” toward the male pathway and promote hub development. Normally, *bab* mutants must be homozygous to see defects in TF formation. However, we found that removing one copy of the *bab1/2* locus (heterozygotes for *bab2AR07* which removes both *bab1* and *bab2* (Couderc et al., 2002)) in a *dsx* mutant background caused a dramatic shift toward hubs over TF (Figs 6G). The hubs seen in this mutant background were positive for two adhesion molecules found in the hub (Fasciclin-3 and N-cadherin), suggesting they represent true hubs rather than just defective TFs. Interestingly, in a few cases we found both a hub and TFs which had formed in the same gonad (Fig. S4C). In an effort to identify a factor similar to *bab* that acts in the hub, we examined *escargot (esg)*, which encodes a transcription factor expressed in those SGPs that form the hub (Gönczy et al., 1992; Le Bras and Van Doren, 2006). However, no shift toward TF/CC formation was observed when we altered *esg* gene dosage in a *dsx* mutant background (data not shown), suggesting that *esg* may not be a principal target for controlling sexually dimorphic niche formation.

**Fig 6.**
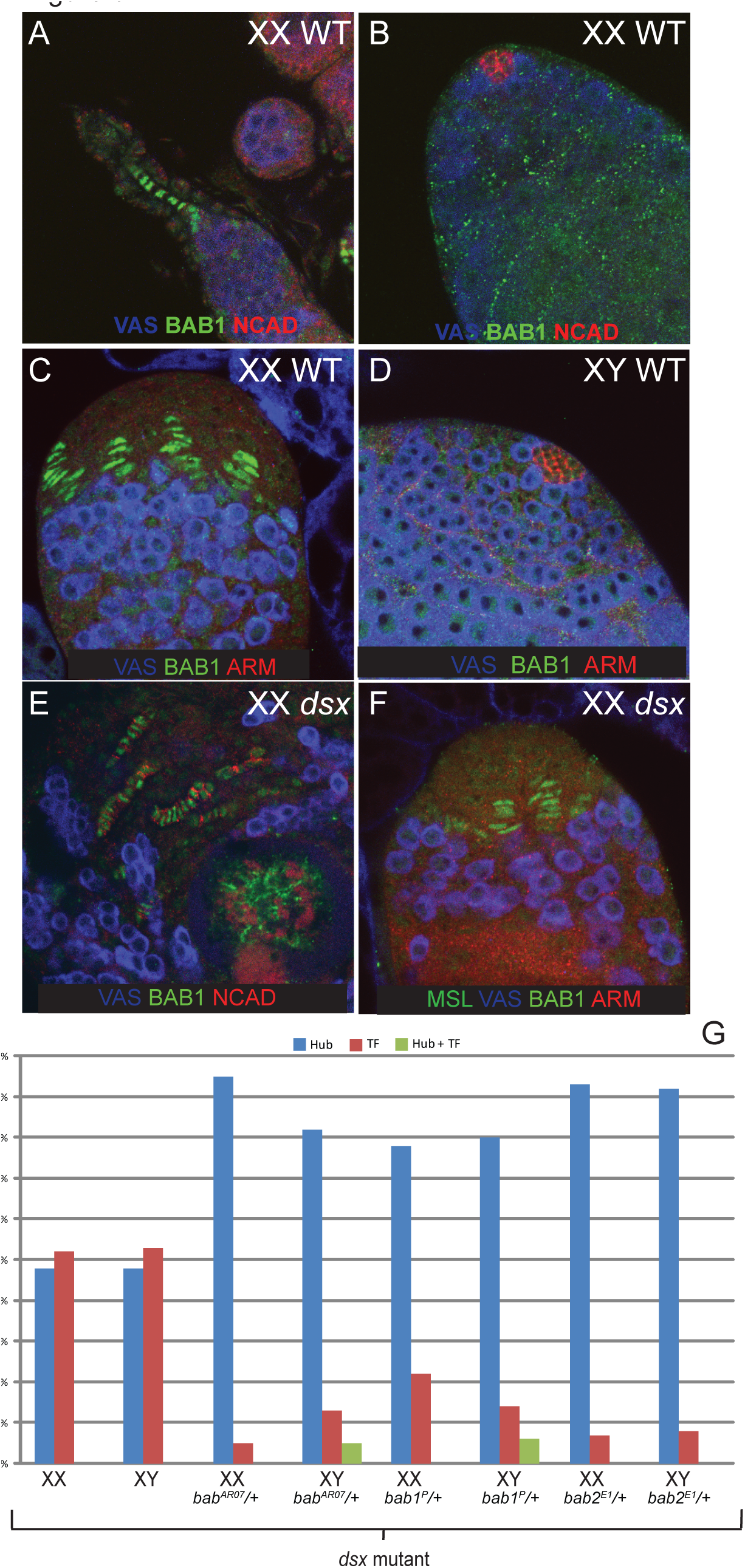
Role of *bab* in sexually dimorphic niche formation. Staining as indicated in all panels. (A,B) BAB1 expression in WT adult female and male. BAB1 is expressed in the female niche, but is absent from the male niche. (C,D) BAB1 expression in WT 3^rd^ instar gonads, present in developing TFs (C), but absent from hubs (D). (E,F) BAB1 expression in terminal filaments on 3^rd^ instar (E) and adult (F) *dsx* mutant gonads. (G) Graph showing genetic interaction with *bab* alleles. *bab2AR07* affects both *bab1* and *bab2, bab1P* affects only *bab1, bab2E1* affects only *bab2* (Couderc et al., 2002). Percentage of gonads with hubs or terminal filaments is indicated. Blue bars indicate hubs, red bars terminal filaments and green bars both hubs and terminal filaments.

*bab1* and *bab2* are thought to be partially redundant, and both act in each tissue for which a role of *bab* has been proposed (Couderc et al., 2002). We observed BAB2 expression in both hubs and TFs (Fig. S5A,B), however BAB1 was found only in the TFs and was absent from hubs (Fig. 6A,B). BAB1 expression begins at the 3^rd^ instar stage, as the female niche is forming ((Godt and Laski, 1995) and Fig 6C), but was absent from the larval hub (Fig. 6D). In addition, an enhancer trap insertion in the first intron of *bab1* recapitulated the sexually dimorphic expression of *bab1* in the female niche, suggesting it’s sex-specific expression is regulated at the level of transcription (data not shown). In *dsx* mutants (both XX and XY), we found BAB1 expression only in the developing terminal filaments, and it was absent from hub cells (Figs 6E,F S5C-H). Thus, in the absence of *dsx*, BAB1 is expressed in female-like niches, but not male-like niches, of both XX and XY gonads, consistent with the stochastic nature of TF/CC vs. hub formation in *dsx* mutants. Finally, we investigated whether it is *bab1* or *bab2* that is important for sexually dimorphic niche development by examining alleles specific to either gene (Couderc et al., 2002). Removing one copy of either gene alone was able to shift the balance towards hubs, similar to that observed with the allele that affects both genes (Figs 6G, S4 D-G). Thus, even though only *bab1* appears to be expressed in a sexually dimorphic manner during niche formation, both *bab1* and *bab2* are required to promote development of the female niche over the male niche.

## DISCUSSION

### *dsx* acts to “tip the balance” between male and female developmental programs

An important question about the creation of sexual dimorphism is how a key transcription factor like DSX regulates a sex-specific developmental program. One model is that DSX could act as a “micromanager” to control many genes required for the formation of sex-specific structures such as the hub or TF/CC. However, we find that these structures can still form in the absence of *dsx* function, but now do so independently of the chromosomal constitution of the animal (XX vs. XY). The hubs and TF/CC that form in *dsx* mutants have many of the characteristics of the wt structures, including the proper morphology and pattern of gene expression, and they can associate with and signal to the germ cells. Therefore, we conclude that DSX does not independently regulate the many different genes that are likely to be required to form these structures. Instead, DSX primarily acts to ensure that the male structures form reliably in XY animals, while the female structures form in XX animals.

DSX is likely to do this by activating male- or female-specific developmental programs that can then function independently of DSX. Thus, DSX would act to “tip the balance” between whether the male (hub) or female (TF/CC) pathway was activated. One way in which DSX might regulate this balance is by influencing the expression of key upstream regulators of these pathways. It was previously known that the *bab* locus was important for TF formation (Sahut-Barnola et al., 1995). Here we show that partial ‘loss of *bab* not only inhibits TF formation in *dsx* mutants (even when only one copy of the *bab* locus is lost), but hubs are present in their place. Thus, *bab* appears to be a key target by which DSX acts to “tip the balance” between male and female development.

Previously, it was shown that *bab* is a direct target of DSX in regulating sex-specific pigmentation in the abdomen and regulates an enhancer in intron 1 of *bab1* (Williams et al., 2008). Interestingly, we find that an enhancer trap in this intron recapitulates sex-specific *bab* expression in the gonad niches, suggesting that *bab* is also directly regulated by DSX in the gonad. Further, our genomic analysis of DSX targets supports the view that this *bab* enhancer is bound by DSX, including in cells of the gonad, and contains evolutionarily conserved consensus DSX binding sites (Clough et al., 2014). The specific enhancer construct that recapitulates sex-specific expression of *bab* in the abdomen (Williams et al., 2008) did not drive expression in the gonad (data not shown). It is likely that regulation of sex-specific gene expression requires both a DSX-responsive element, and elements that control tissue-specificity, as has been reported for the *Yp1/2* locus (Coschigano and Wensink, 1993). Thus, the *bab* sex-specific abdomen enhancer may respond to DSX but lack the sequences necessary for driving expression in the gonad. If *bab1* is a direct DSX target, it is somewhat surprising that BAB1 expression is stochastic in *dsx* mutants; on in developing TFs and off in hubs. We might have expected that a DSX target would show intermediate expression in the absence of DSX, with a loss of repression by DSX^F^ and loss of activation by DSX^M^. It is possible that either autoregulation by the BAB proteins, or cross-regulation by an additional upstream DSX target, leads to an “on/off” pattern of BAB1 expression as opposed to intermediate levels.

These data indicate that DSX influences the sexually dimorphic development of two different tissues (abdomen and gonad), with dissimilar developmental programs, by regulating the same downstream target locus, *bab*. Interestingly, sexually dimorphic pigmentation in *Drosophila* is a relatively recent and rapidly evolving trait, and sex-specific regulation of *bab* in the abdomen unexpectedly persists in species that do not exhibit sexually dimorphic abdominal pigmentation (Kopp et al., 2000). Since sexual dimorphism in the gonad is more highly-conserved among animal species, it may be that the regulation of *bab* by DSX in the gonad is the more ancient role for the *dsx-bab* regulatory interaction, and that this has subsequently been co-opted to control species-specific sexual traits, such as pigmentation, that evolve more rapidly.

### The timing of *dsx* action in gonad sex determination

We have found evidence for both early and late roles for *dsx* in promoting gonad sexual dimorphism. In *dsx* mutants, all sexually dimorphic cell types that we have examined initiate the male pattern of development in both XX and XY embryos, including the pigment cell precursors, msSGPs and hub cells (DeFalco et al., 2003; Le Bras and Van Doren, 2006; DeFalco et al., 2008). This would seem to indicate that the primary role of *dsx* in the early gonad is to prevent the male program of development in females. However, we show here that the sex-specific pattern of SGP gene expression is intermediate, between male and female, in *dsx* mutants (Fig 3C) why would an intermediate level of sex-specific gene expression give rise to an initiation of the male pathway? It is relatively easy to resolve this paradox for the pigment cell precursors and msSGPs since their development is regulated in a non-autonomous manner by signals from the SGPs (DeFalco et al., 2008). It may be that an intermediate level of a male-specific signal (e.g. Wnt2 for induction of pigment cell precursors (DeFalco et al., 2008)) is sufficient to induce the male pattern of development.

However, this is more difficult to explain for hub cell formation, where we have shown that the sex determination pathway acts at least partly in a cell-autonomous manner to control hub development (Le Bras and Van Doren, 2006). In mosaic gonads with a mixture of male and female SGPs, the hub is formed only from male SGPs (Le Bras and Van Doren, 2006). Therefore, why would cells with an intersexual pattern of gene expression follow the male pathway initially? Part of the explanation may lie in the normal difference in timing between hub development, which occurs at the end of embryogenesis, and TF/CC formation, which occurs in late 3^rd^ instar larvae. It may be that, even though both male and female gene expression is initiated in *dsx* mutant SGPs, this is sufficient to induce hub formation initially, since this normally occurs earlier than the female pathway of TF/CC development. This idea is supported by our finding that *dsx* mutant hubs are somewhat different than normal “male” hubs, since *dsx* mutant hubs are able to switch to TF/CC at the time that these female structures normally form. This is in contrast to what is observed when hubs form under conditions where DSX^M^ is expressed (*tra2ts* shifted from male to female, Fig 4E). These hubs are clearly resistant to switching to TF/CC formation even when switched to DSX^F^ expression prior to the time when TF/CC would normally form. Thus, *dsx* mutant hubs retain a greater degree of sexual plasticity than do wt male hubs.

A different result is found when gonads develop first under control of DSX^F^, and are then switched to DSX^M^. SGPs that have a female identity early are still fully capable of forming a hub after switching to DSX^M^ expression as second instar larvae, even though this is past the time when a hub would normally form. Thus, the female mode of development is not “locked in” as early as the male mode of development. In contrast, when DSX^F^ expression is maintained throughout the normal time window of TF/CC formation (later L3 larvae), these structures are now maintained even if DSX^M^ is expressed later. Thus, the requirement for the male vs. female forms of DSX in gonad niche development reflects the timing of when these niches normally form. It is possible that DSX may have later roles in maintaining the proper function of these structures, similar to the role we see for DSX in maintaining the pigment cell precursors in older gonads (Fig 5C,D). However, later niche function is difficult to assess with this assay due to the requirement that the sex of the germline must match the sex of the soma for proper gametogenesis.

### *dsx* likely acts on one primordium to create male vs. female niches

Another important question for the creation of sexual dimorphism is in which cells does the sex determination pathway act to influence male vs. female development? Further, does it act in one primordium to allow it to develop into either male or female structures, or do the male and female structures come from distinct primordia? For example in Drosophila, the male and female reproductive tracts are predominantly formed from distinct male and female primordia within the genital disc; only one of these primordia will grow in either sex to give rise to the majority of the reproductive tract, while the repressed primordia makes a minor contribution (Estrada et al., 2003). In contrast, our data support a model where the hub and TF/CC come from the same primordium, and *dsx* acts on these cells to allow them to contribute to either the male or female gonad niche. It has been shown that the hub is formed from anterior SGPs of the embryonic gonad (Le Bras and Van Doren, 2006). It has also been proposed that TF/CC arise from anterior SGPs in the female (Asaoka and Lin, 2004). Further, the data we present here suggests that, in *dsx* mutants, cells that have taken on hub identity can revert and form TF/CC. This clearly supports a model where these key cell types in the male and female gonad niches are closely related to one another and represent alternate developmental paths of the same primordium.

It is interesting we do not usually find hubs and TF/CC forming in the same gonad in *dsx* mutants. This indicates that the cells of the hub-TF/CC primordium are coming to a “group decision” about what their sex should be and which type of niche to form. One might have imagined that, in the absence of *dsx* function, each cell might make a cell-autonomous choice of what sex to be, and whether to take on hub or TF/CC identity. Under this model, some cells could follow the hub developmental program, while others could form TF and CC. This is clearly not the case. Thus, there must be some non-autonomous component to the choice of male vs. female identity in the primordium, such that all cells follow the same developmental path. In a few cases, evidence for both hubs and TF forming in the same gonad was observed when *bab* was compromised in a *dsx* mutant background (Fig. 6G). This may indicate that the signal governing the “group decision” was affected in this genotype, suggesting that it may be downstream of factors such as *bab*. As mentioned above, our previous work revealed that when a gonad consists of a mosaic of male and female cells, female cells are excluded from the hub (Le Bras and Van Doren, 2006). Thus, our evidence indicates that the choice between hub and TF/CC identity is regulated both cell-autonomously, dependent on a cell’s endogenous sex determination information, as well as non-autonomously, via signaling between cells of the gonad niche primordium.

### Parallels between flies and mammals

The model that is emerging for how *dsx* regulates sexual development of the Drosophila gonad is strikingly similar to how sex determination is regulated in the mammalian gonad. Both utilize a combination of autonomous and non-autonomous factors to decide their sexual identity and activate the male or female developmental program. When mouse gonads are mosaic for XX and XY cells, a testis can form in the presence of sufficient XY cells (Burgoyne et al., 1988). The only cells of the testis that show a bias toward being XY are the male Sertoli cells. However, even XX cells can form Sertoli cells if intermingled with enough XY cells (Palmer and Burgoyne, 1991). Thus, there is an autonomous contribution of a cell’s sex chromosome constitution to male Sertoli cell formation, but also a non-autonomous agreement among the cells about whether to be male or female, and even XX cells can take on a male identity. We now know that the autonomous signal is based on expression of the Y chromosome gene SRY, but that the regulation of genes acting downstream of SRY, such as SOX9, is influenced by non-autonomous signals, namely FGF9 in males and WNT4 in females (DiNapoli and Capel, 2008). In this way, a balance of autonomous and non-autonomous information controls sexual identity in the primary somatic cells that support the germline in the mouse, as we have shown in Drosophila for the niches that support the germline stem cells.

Another parallel between flies and mammals lies in the sexual pasticity of gonadal cell types, such as the switching between hubs and TFs we observe here. Male cyst cells of the testis have also demonstrated plasticity, transforming into female follicle-like cells in the absence of the transcription factor *chinmo* (Ma et al., 2014, Grmai et al., 2018). Such plasticity is also observed in the mouse gonad. Male gonadal structures can initially form in mice mutant for the DSX homolog DMRT1 (Raymond et al., 2000), but they are not properly maintained, and eventually revert to more female identity (Sertoli cell to Granulosa cell, Matson et al., 2011). Similarly, deletion of the female-promoting factor FOXL2 can induce the opposite transformation (Uhlenhaut et al., 2009). One interesting difference between flies and mice is that mouse gonads appear sexually labile even after birth, with mutation of DMRT1 or FOXL2 able to cause sex reversal by controlled, postnatal deletion (Uhlenhaut et al., 2009; Matson et al., 2011). In contrast, flies appear to have a more defined developmental time window for sexual plasticity, which depends on when the male or female structures normally develop. However, such sexual transformations emphasize an important aspect by which the sex determination pathway regulates sexual dimorphism; male and female cell types may appear dramatically different, but are often highly analogous, and can even transform from male to female and vice versa, indicating that sex determination need only alter relatively specific aspects of otherwise similar cell types.

## MATERIALS AND METHODS

### Fly Stocks

The following stocks were used: *dsx^1^, Df(3R)dsx^3^, upd-Gal4* (T. Xie), *y^1^w^1118^, P{w^+mc^=UAS-Gal4.H}12B, w^1118^; P{w^+mc^=UAS-GFP.nls}14, Dp(1:Y) B^S^, bw^1^ tra2^ts1^, y^1^/Dp(1;Y) B^S^, cn^1^ tra2^ts2^ bw, dad-lacZ, esg^M5-4^* (S. DiNardo), *hedgehog-lacZ* (A. Spradling), *esg^G66B^* (Kassis Lab), *bab2^ARO7^* (F. Laski), *bab2^E1^* (F. Laski) *bab1^P^ cn8 ry ca* (F. Laski), *w^*^; P{w^+mW.hs^=GawB] bab1^Agal4-5^, w^*^; P{w^+mW.hs^=GawB] bab1^Pgal4-2^, w^-^;; bab1-Gal4e* (A. Gonzales-Reyes), *S3aG bab1 sub1.6 WT* (S. Carroll), *S3aG bab1 sub1.6 Dsx1,2 KO* (S. Carroll), *BPSMG2 bab1 #1* (S. Carroll). *w*^1118^ is the wild-type control. Unspecified stocks can be found on Flybase (http://flybase.bio.indiana.edu).

### Antibody Stainings and In Situ Hybridization

Adult testes and ovaries were dissected in PBS and fixed 30 minutes at RT in 4.5% formaldehyde in PBS containing 0.1% Triton X-100 (PBTx). Staining was as described {Gonczy, 1997 #100} and samples were mounted in 2.5% DABCO.

The following antibodies (sources) were used: chicken anti-VASA (K.Howard) at 1:10,000; rabbit anti-VASA (R. Lehmann) at 1:10,000; rat anti-VASA (Developmental Studies Hybridoma Bank, DSHB, A.C. Spradling/D. Williams) at 1:50; rabbit anti-SOX100B at 1:1,000 (S. Russell); mouse anti-EYA 10H6 (DSHB, S. Benzer/N.M. Bonini) at 1:25; rabbit anti-GFP (Torrey Pines) at 1:2,000; mouse anti-GFP (Santa Cruz) at 1:50; mouse anti-FAS3 7G10 (DSHB, C. Goodman) at 1:30; rabbit anti-β-GAL (Cappel) at 1:10,000; mouse anti-β-GAL (Promega) at 1:10,000; rat anti-DN-cadherin Ex #8 (DSHB, T. Uemura) at 1:20; guinea pig anti-TJ (D. Godt) at 1:3000; rabbit anti-msl2 (M. Kuroda) at 1:1,000, rabbit anti-Zfh-1 (R. Lehmann) at 1:5,000, rabbit anti-STAT92E (S. Hou) at 1:1,000, mouse anti-LaminC LC28.26 (DSHB, P.A. Fisher) at 1:20, mouse anti-Engrailed 4D9 (DSHB, C. Goodman) at 1:2, rabbit anti-phospho histone H3 (Upstate) at 1:5,000, rabbit anti-BAB1 (S. Carroll), rat anti-BAB2 R10 (F. Laski), mouse anti-armadillo N2 7A1 (DSHB, E. Wieschaus). The following secondary antibodies were all used at 1:500: Cy5 goat anti-chicken (Rockland), Alexa 546 goat anti-chicken, Alexa 546, 488 or 633anti-rabbit, Alexa 546, 488 or 633 goat anti-mouse. Alexa antibodies are from Molecular Probes (Invitrogen, Carlsbad, CA, USA).

In situ hybridization performed as described (DeFalco et al., 2003), with a colorimetric (NBT/BCIP) substrate, with antibody staining performed subsequently to determine genotype.

### Genotyping and Sexing

*GFP*-expressing balancer chromosomes were used to identify homozygous mutant animals. Sex chromosome genotype was determined using *lacZ*-expressing X chromosomes (DeFalco et al., 2003), Y chromosomes marked with B^S^, or using a male-specific anti-msl1 (M. Kuroda).

### *tra2* heat shifting experiments

*tra2^t1^* males were mated to *tra2^ts2^* virgins at 18°C or 29°C. Animals were raised and temperatures shifted at either the 2^nd^ instar or pupal time points. Gonads were dissected at the 3^rd^ instar or 1day old adults.

## ACKNOWLEDGEMENTS

We are very grateful to members of the fly community that supplied us with fly stocks and reagents for this work, as specifically cited in Materials and Methods. We also thank the Bloomington Stock Center (Indiana University), the Developmental Studies Hybridoma Bank (University of Iowa), and the Drosophila Genome Resource Center for reagents. In addition, we would like to thank members of the Van Doren Lab for helpful discussions. Imaging was performed at the Integrated Imaging Center at The Johns Hopkins University. This work was supported by the National Institutes of Health (GM084356 and GM113001).

---

**Supp Fig 1.**
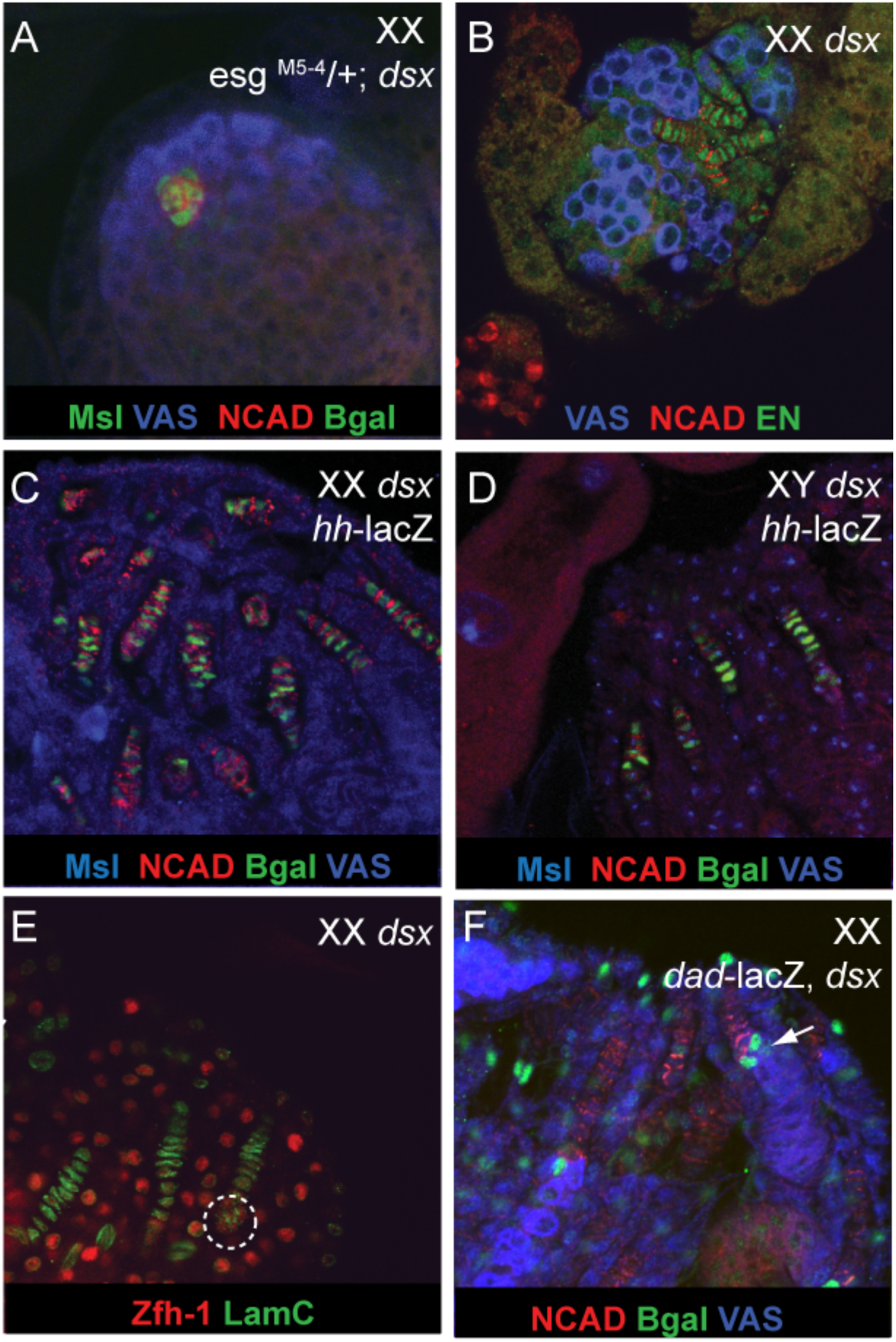
Staining as indicated in all panels. (A) *esg* enhancer trap expressed in the hub of XX *dsx* mutant adult. (B) Engrailed expression in terminal filament of XX *dsx* mutant adult. (C,D) *hedgehog-* lacZ in XX and XY *dsx* mutant terminal filaments. (E) Cap cells in XX *dsx* mutant adult with terminal filaments (cap cells are co-labeled for LamC and Zfh-1). (F) dad-lacZ in germ cells adjacent to the terminal filaments in *dsx* mutant gonads.

**Supp Fig 2.**
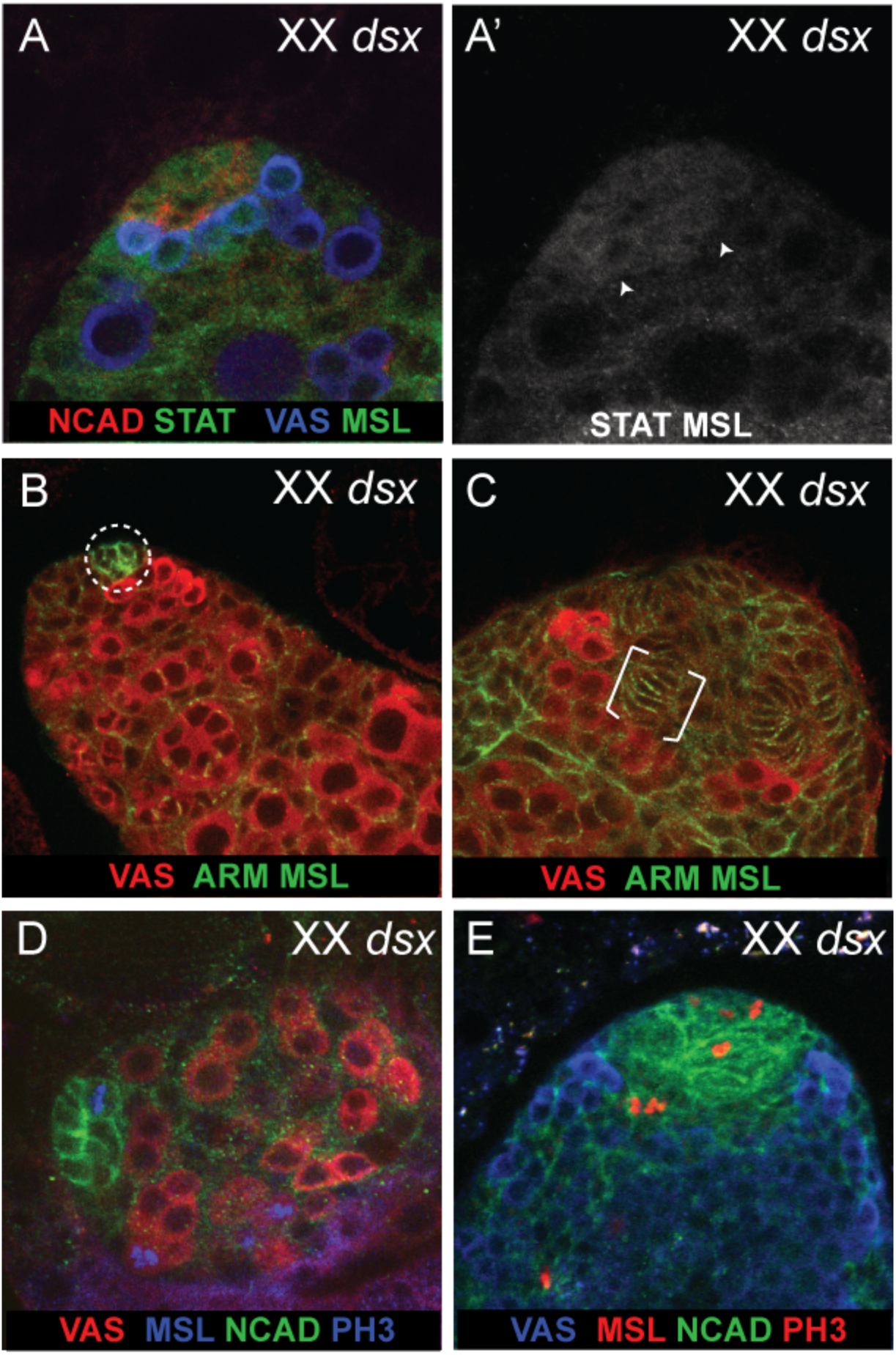
Staining as indicated in all panels. (A) STAT92E localization in germ cells adjacent to a hub in *dsx* mutants. (B) XX *dsx* mutant 3^rd^ instar gonad that retains hub morphology. (C) Developing terminal filaments in an XX *dsx* mutant 3^rd^ instar gonad. (D) Proliferation in XX *dsx* mutant 2^nd^ instar hub. (E) Proliferation in XX *dsx* mutant 3^rd^ instar developing TFs.

**Supp Fig 3.**
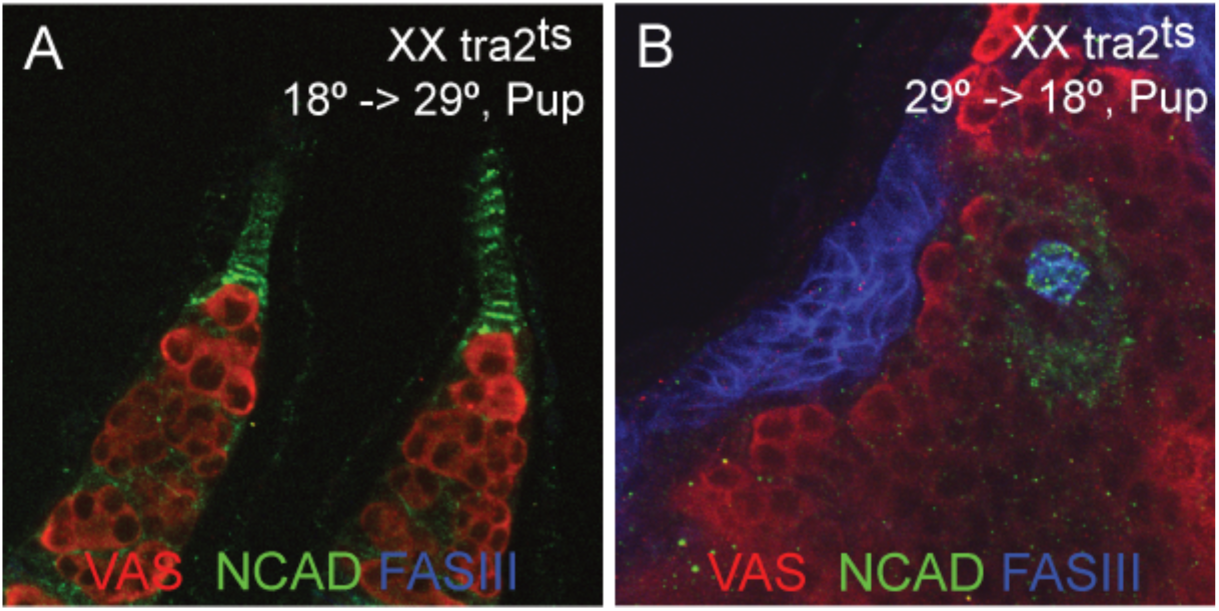
Staining as indicated in all panels. (A) XX tra2^ts^ shifted from 18 to 29 at pupal stages develops TFs. (B) XX tra2^ts^ shifted from 29 to 18 at pupal stages develop hubs.

**Supp Fig 4.**
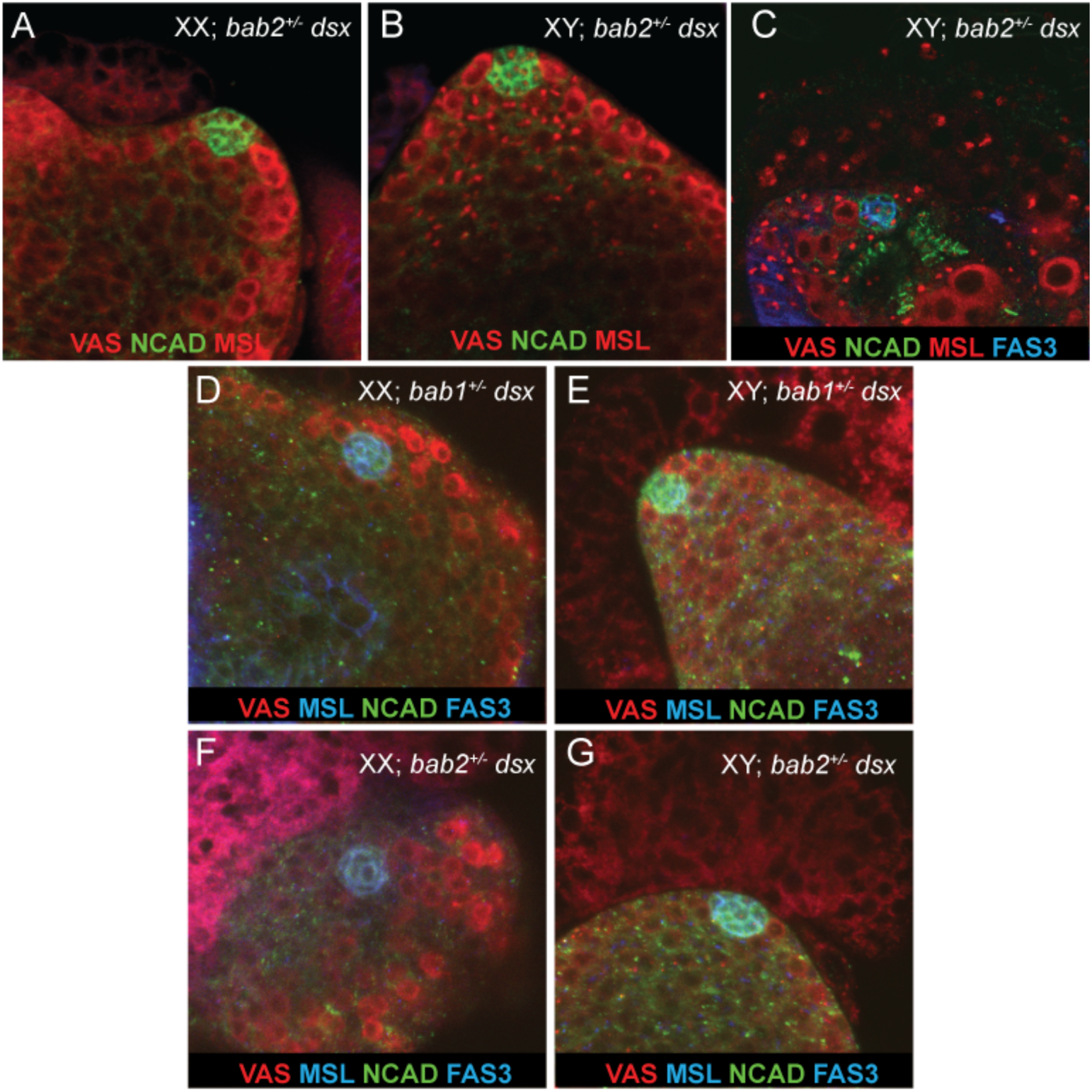
Staining as indicated in all panels. (A-C) Gonads from *dsx* mutants missing one copy of *bab2^AR07^*. (C) Example of a gonad containing both a TF and hub. (D,E) Gonads from *dsx* mutants missing one copy of *bab1^P^*. (F, G) Gonads from *dsx* mutants missing one copy of *bab2^E1^*. Anti-MSL2 staining appears as punctuate nuclear staining in the soma of XY gonads.

**Supp Fig 5.**
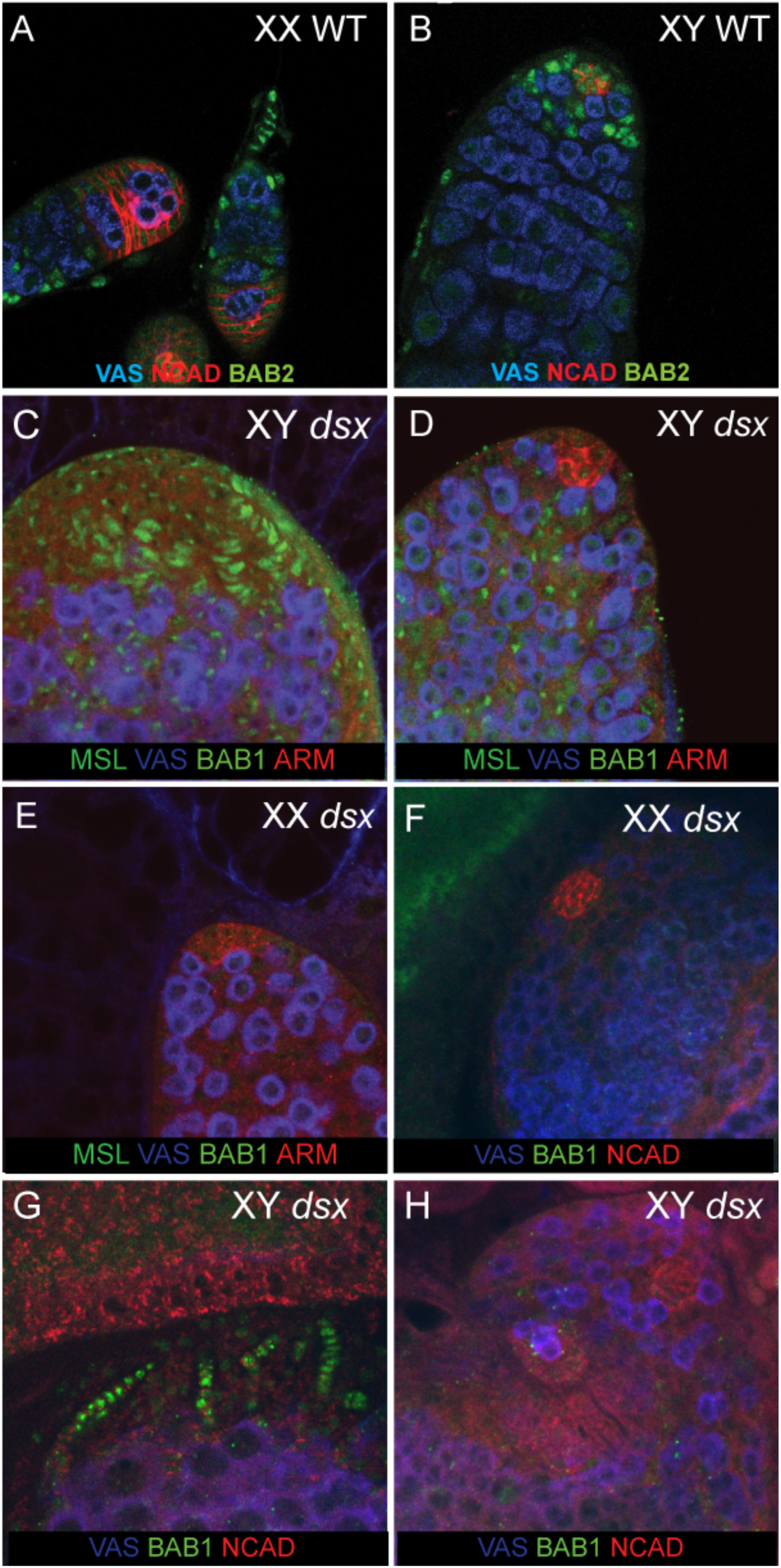
Staining as indicated in all panels. (A,B) BAB2 staining in wild type adult gonads. BAB2 is present in both terminal filaments (A) and hubs (B). (C,D) BAB1 is present in XY 3^rd^ instar *dsx* mutant gonads in developing terminal filaments (C), but is absent from the hub (D). (E) BAB1 is absent from XX *dsx* 3^rd^ instar hubs. (F) BAB1 is absent from XX *dsx* mutant adult hubs. (G,H) BAB1 is present in XY *dsx* mutant adult TFs (G), but absent from adult hubs (H).

